# Antibodies to ILT3 abrogate myeloid immunosuppression and enable tumor killing

**DOI:** 10.1101/2021.12.18.472552

**Authors:** Philip E. Brandish, Anthony Palmieri, Gulesi Ayanoglu, Jeanne Baker, Raphael Bueno, Alan Byford, Michael Caniga, Craig Chappell, Holly Cherwinski, Daniel J. Cua, Xiaoyan Du, Laurence Fayadat-Dilman, Brian E. Hall, Hassan Issafras, Barbara Joyce-Shaikh, Veronica Juan, Rachel D. Levy, Andrey Loboda, Jared Lunceford, Carl Mieczkowski, Michael Meehl, Yujie Qu, Michael Rosenzweig, Latika Singh, Peter Stivers, Peter J. Tramontozzi, Kimberly Vermilya, Karin Vroom, Douglas C. Wilson, Chungsheng Zhang, Jie Zhang-Hoover

## Abstract

Tumor myeloid suppressor cells impede response to T cell checkpoint immunotherapy. Immunoglobulin-like transcript 3 (ILT3, gene name, *LILRB4*) expressed on dendritic cells (DCs) promotes antigen-specific tolerance. Circulating monocytic MDSCs that express ILT3 have been linked to clinical outcomes and a soluble form of ILT3 is elevated in certain cancers. We find that *LILRB4* expression is correlated with Gene Expression Profile of T-cell inflamed tumor microenvironment shown to be significantly associated with response to the anti-PD1 antibody pembrolizumab across several tumor types. A potent and selective anti-ILT3 mAb effectively antagonized IL-10 polarization of DCs and enabled T cell priming. In an MLR assay anti-ILT3 combined with pembrolizumab afforded greater CD8^+^ T cell activation compared to either agent alone. Anti-ILT3 antibodies impaired the acquisition of a suppressive phenotype of monocytes co-cultured with SK-MEL-5 cancer cells, accompanied by a reduction in surface detection of peptidase inhibitor 16, a *cis* interaction candidate for ILT3. Growth of myeloid cell-abundant SK-MEL-5 tumors was abrogated by ILT3 blockade and remodeling of the immune tumor microenvironment was evident by CyTOF. These data support the testing of anti-ILT3 antibodies for the treatment of a wide range of solid tumors replete with myeloid cells.

## INTRODUCTION

New therapeutics targeting inhibition of the cytolytic T cell response such as PD1-blocking antibodies have increased overall survival in a range of cancers either alone or in combination with chemotherapy, targeted therapy, or other immune agents. At the same time, a great deal of unmet medical need remains and there are large gaps in our knowledge of mechanisms at work when tumors resist treatment (1). Not least among the myriad of mechanisms of resistance is myeloid cell-driven tolerance and immune suppression (2-5). Myeloid cells are both diverse and plastic with respect to phenotype and function and the complexity of this compartment in the tumor microenvironment has come under scrutiny with the application of high dimensional profiling technologies (6-10). In the clinical oncology setting, myeloid cells have gained attention in the context of activation of antigen-presenting cells (dendritic cells and macrophages) as integral to the anti-tumor immune response, for example via activation of innate nucleic acid sensing pathways (11). In addition, attention has focused on a broad class of immature myeloid cells that functionally suppress T cell activity termed myeloid-derived suppressor cells (MDSCs) (5, 12, 13). To date though, targeting of the myeloid compartment has not yielded approved new therapies.

We and others have been intrigued by the underlying biological analogies between the body’s response to cancer and two important homeostatic processes, specifically immune tolerance and tissue injury and repair. Tolerance to self-antigens and acquisition of tolerance in the periphery, when appropriate, is essential to avoid rampant auto-immunity, but also is highly relevant at the maternal-fetal interface and in the setting of organ transplant (14-18). Tissue injury and repair, the wound healing response, is characterized by an orchestrated management of the immune response, including infiltration with immune suppressive myeloid cells (19, 20). The analogies to cancer and the potential for normal processes of self-tolerance and wound healing to enable the continued growth of tumors were striking to us. In both tolerance and immune suppression, a family of surface receptors, called the ILTs or LILRs, have been noted for their roles. In particular ILT3 gained our attention at the intersection of several observations: (1) Studies using organ transplant or mixed lymphocyte culture showed that ILT3 expression on antigen-presenting cells is associated with antigen-specific tolerance and suppression of T cell activation (21-25). (2) Myeloid-derived suppressor cells (MDSCs) are recognized as critical players in the tumor microenvironment (5, 12, 13, 26). The monocytic variety express ILT3 and the level of ILT3 expression on circulating MDSCs has been connected to survival in patients with lung cancer (27). In patients with previously treated metastatic bladder cancer, high baseline circulating monocytic MDSC count was associated with a dramatically lower overall survival after treatment with nivolumab compared to patients with a low MDSC count (28). (3) Soluble ILT3 levels were reported to be elevated in patients with certain types of cancer and was reported to be immunosuppressive (25, 29, 30). (4) Identification of ILT3 as emblematic of a myeloid cell signature associated with T cell inflammation in solid tumors (this work).

ILTs can be either activating or inhibitory depending on whether their intracellular domains contain immunoreceptor tyrosine-based activating (ITAM) or immunoreceptor tyrosine-based inhibitory (ITIM) motifs (31). Given the presence of ITIM motif in the ILT3 sequence and observations of inhibitory activity upon ILT3 co-ligation in vitro (32, 33), ILT3 likely functions as an inhibitory receptor.

ILT3 is expressed on monocytic myeloid cells such as myeloid-derived suppressor cells (MDSCs), monocytes, macrophages, dendritic cells, but not granulocytes (31). More recently, Deng and co-workers identified a clear role for ILT3 in acute myeloid leukemia (AML) cells, characterized signaling pathways that drove T cell suppression, and provided evidence that blocking ILT3 might be beneficial in the treatment of AML (34, 35). ILT3 was found to bind to CD166 and ApoE in vitro, while in vivo functional significance remains to be determined (34-36). In the work reported here we explored the potential for ILT3 as a drug target for intervention in solid tumors where myeloid cells drive immune suppression or erroneous treatment of the tumor as self.

## RESULTS

### Molecular and plasma protein data suggest a role for ILT3 in human cancer

We analyzed the expression of the gene for ILT3, LILRB4, in two contexts. First, in the public TCGA data set, we found that LILRB4 is co-expressed and highly correlated with the 18-gene T cell-inflamed gene expression profile (GEP) signal that is correlated with the likelihood of a response to the anti-PD1 antibody pembrolizumab (37) (Fig. 1A). LILRB4 expression was not associated with mutational load. We found essentially the identical pattern in the Merck & Co., Inc., Kenilworth, NJ, USA-Moffitt data set (not shown). As mentioned, GEP signature high tumors are more likely to respond to pembrolizumab than GEP signature low tumors and those characteristics are illustrated for context in Fig. 1B taken from a recent report by some of us (38). Extending the findings from analysis of TCGA data, analysis of LILRB4 expression in samples collected during clinical trials of pembrolizumab (39) confirmed significant correlation with GEP and even more significant correlation with mMDSC signature (Fig. 1C). In this pan-cancer dataset, LILRB4 showed 0.67 correlation with GEP which had highly significant association with response to pembrolizumab (AUC=0.63; p-value <<0.0001) (39). LILRB4 had even higher correlation (0.79), with the consensus signature of monocytic MDSC developed at Merck & Co., Inc., Kenilworth, NJ, USA, (39) which showed highly significant association with resistance to pembrolizumab in GEP adjusted analysis (AUC=0.56; p-values=0.0001). These data suggest that ILT3 is linked to the biology underlying responsiveness to pembrolizumab but we cannot from these data alone infer causality or direction of effect. Given the prior knowledge of ILT3 biology that is available, it is reasonable to hypothesize that inhibiting ILT3 would be beneficial (rather than activating ILT3).

**Figure 1:**
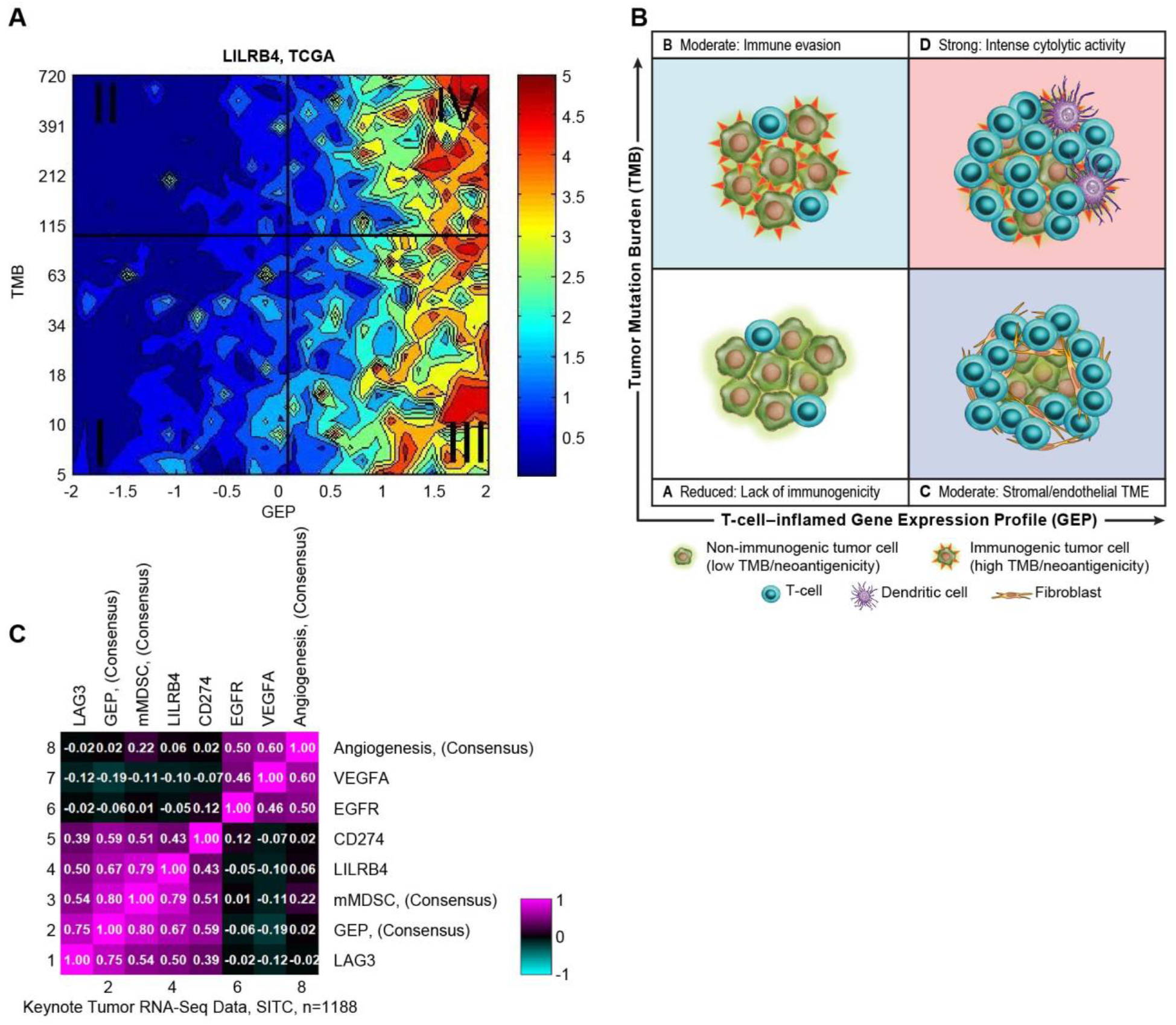
Expression of LILRB4 as a function of GEP and TMB and association with clinical response to pembrolizumab. (A) Tumor Mutational Burden (TMB) is plotted against the 18-gene GEP score for tumor samples in The Cancer Genome Atlas (TCGA) data set. The contour plot illustrates the association of LILRB4 with TMB and GEP. Blue and red represent under- and overexpression, respectively. LILRB4 is highly co-expressed with GEP regardless of mutational burden. TMB cut-off was set at 100 and GEP cut-off corresponds to 55th percentile value for pan-cancer cohort. **(B)** Illustration of characteristics of tumors within the quadrants identified in the left panel plot as described in Critescu et al. and reproduced with permission from Science. **(C)** Covariance patterns of gene signatures derived from RNA-seq data from pembrolizumab clinical trial samples. Expression of LILRB4 is highly correlated to both GEP and monocytic MDSC signature.

Turning to protein expression, we analyzed by flow cytometry the presence of ILT3 protein on the surface of tumor-resident or infiltrating immune cell populations in several common solid tumor types (Fig. 2A). The published literature describes ILT3 as restricted to the monocytic myeloid lineage (31, 32, 40) and that is essentially what we have found both in human tumors (Fig. 2A and Fig. S1A) and in human SK-MEL-5 tumors grown in humanized mice (Fig. S1B). We also measured soluble ILT3 levels in the plasma of patients with various types of cancer using a fully validated assay developed at Merck & Co., Inc., Kenilworth, NJ, USA (Fig. 2B). We found higher levels in some cancers but not others, although in no case was the difference more than 3-or 4-fold on average. We made these measurements on the basis of prior work reporting elevated levels in cancer patient plasma (25). The same report indicated the presence of an alternative LILRB4 transcript accounting for at least some fraction of the soluble material, but we were not able to corroborate those findings (data not shown). Nevertheless, the elevation of soluble ILT3 in cancer patient plasma is a reproducible and potentially etiologically meaningful finding.

**Figure 2.**
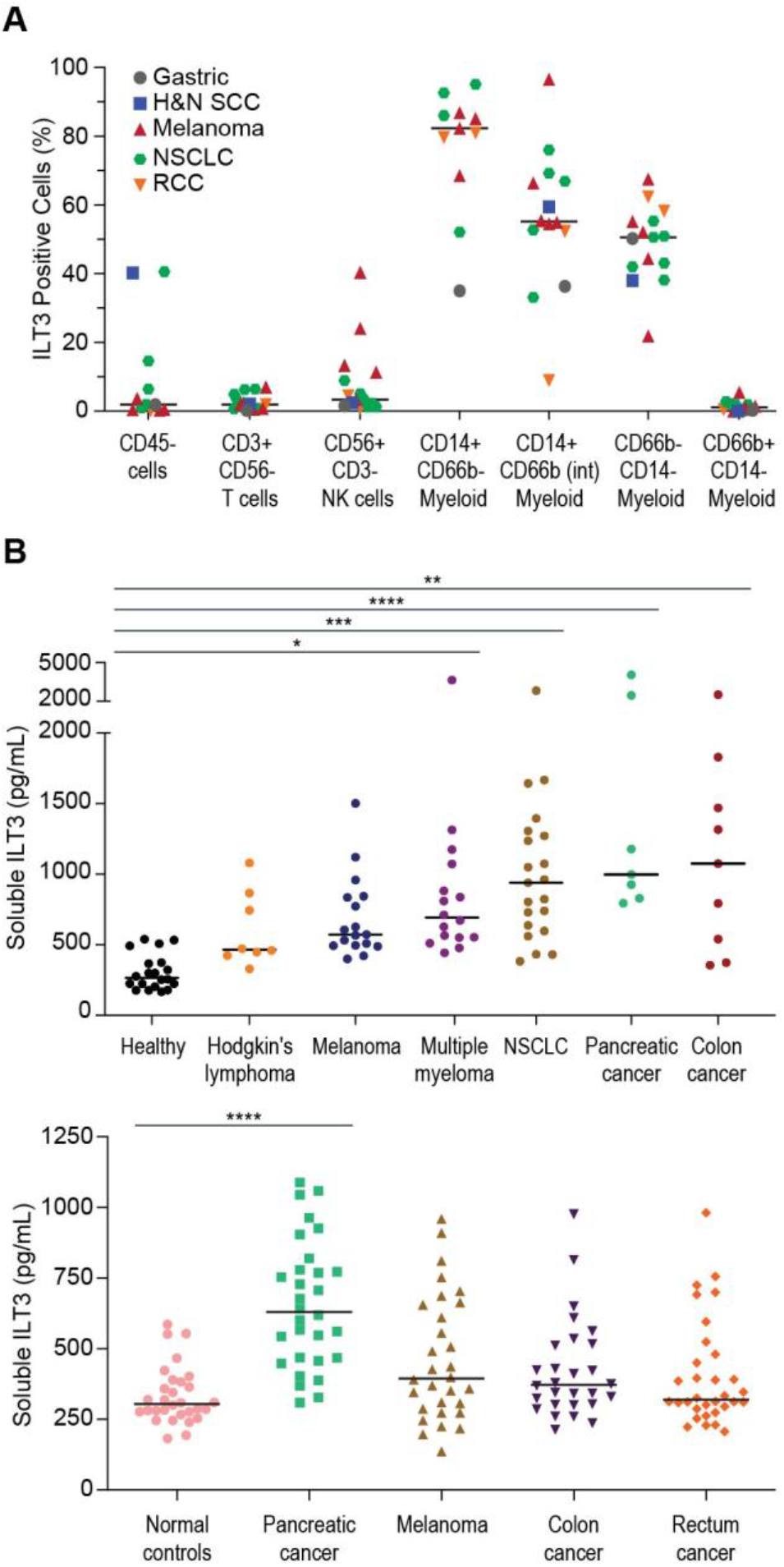
Characterization of ILT3 expression. (A) Surface expression of ILT3 was measured on immune cells from human tumors. Tumor tissue was procured from a third party vendor and processed for flow cytometry as described in the Materials and Methods section. Anti-ILT3 mAb clone ZM4.1 was used to assess ILT3 expression as frequencies of ILT3-positive cells over isotype controls using conventional flow cytometry. Shown are the frequencies of ILT3 positive cells within CD45-cells (tumor and tumor stroma), tumor infiltrating immune cells (CD3+ CD56-T cells, CD56+ CD3-NK cells), CD14+ CD66b-myeloid cells, CD14+ CD66bINT myeloid cells, CD66b-CD14-myeloid cells, and CD66b+ CD14-myeloid cells from 15 tumor tissue donors. The 15 tumor samples are 1 gastric tumor sample, 1 head and neck squamous cell carcinoma (HNSCC) sample, 5 melanoma samples, 6 non-small cell lung carcinoma (NSCLC) samples, and 2 renal cell carcinoma (RCC) samples. Individual symbols represent individual values and horizontal lines indicate group medians. (B) Soluble ILT3 levels were measured in human plasma. Samples were obtained from Bioreclamation (left) and Sanguine (right) and soluble ILT3 levels were determined using a fully validated custom sandwich ELISA assay developed at Merck & Co., Inc., Kenilworth, NJ, USA. ANOVA and follow up with Dunnet’s test found significantly elevated levels of soluble ILT3 compared to control in Multiple Myelomoa, NSCLC, Pancreatic cancer and Colon cancer in the Bioreclamation set and in Pancreatic Cancer in the Sanguine set, *p val 0.0332, ** p val 0.0021, *** p val 0.0002, **** p val < 0.0001.

### Creation of highly selective antibodies as tools to probe the function of ILT3

In concert with the published literature the gene and protein expression data presented above provided an impetus for us to develop high quality pharmacologic tools with which to probe the biology of ILT3 in relevant functional models. Initial antibody campaigns to generate research tools and characterization of other clones in the published or patent literature (e.g. TolerX clone 9B11, Patent # US 8,901,281 B2) taught us quickly that a great majority of clones would not be selective for ILT3 over other proteins in the ILT / LILR family likely due to high sequence similarity within the family (41). The primate ILT / LILR family is generally recognized to be evolutionarily divergent from the rodent orthologs (31). Therefore, we prioritized a classical rodent protein antigen immunization approach to generate selective antibodies in part on the speculation that a rodent campaign would bias toward epitopes not conserved across species and therefore be more likely to target regions related to the divergence of function and potentially ligands, loosely interpreted as the functionally relevant surfaces.

To enable testing for specificity we created a panel of stable CHO cell lines expressing individually the main ILT family members (ILT2, ILT3(D223), ILT3(G223), ILT4, ILT5, ILT7, ILT8, ILT11, LILRA1, LILRA2, and LILRB5) as well as rhesus ILT3 and used these to screen parental clones from mouse and rat immunizations (data not shown). The utility of this approach is illustrated in Fig. 3 – data are shown for a mouse anti-human antibody, labeled clone 52B8. Epitope binning, consideration of expression yield and quality data, as well as review of amino acid sequences led us to prioritize four clones, 52B8, 51H11, 42F6 and 22A5 for further study as recombinant chimeric human IgG4s (Table 1 and data not shown).

**Table 1.** Properties of key clones arising from rodent anti-human ILT3 antibody campaigns.

| Clone | Epitope Bin | Binding Affinity for Human ILT3 Monovalent Kd (nM) | Cell ELISA Human ILT3 EC<sub>50</sub> (ng/ml) | Human DC-TNFα Assay EC<sub>50</sub> (ng/ml) | PK - C<sub>max</sub>, C<sub>trough</sub> 3 mpk i.p. (μg/ml) |
| --- | --- | --- | --- | --- | --- |
| 52B8 | 1 | 0.46 | 15.5 | 1.9 | 45.7, 17.8 |
| 51H11 | 2 | 6.80 | 38.4 | active | 23, 5 |
| 42F6 | 3 | 0.99 | 50.5 | 4.6 | 27, 11.5 |
| 22A5 | 4 | 0.76 | 37.4 | 9.1 | 31.4, 7 |

Note: Data are shown for chimeric human IgG4(S228P) antibodies, that is, rat or mouse variable domains grafted onto a human IgG4 framework to enable direct comparisons. Cross-blocking experiments established that these antibodies bind to distinct epitopes: The epitope bin numbers are arbitrary.

**Figure 3.**
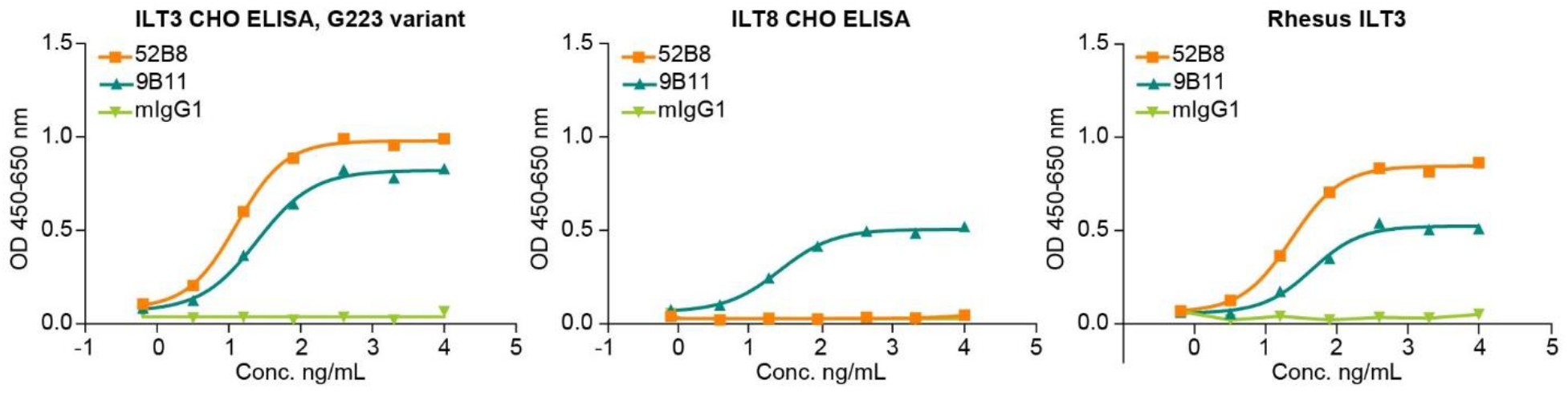
Selectivity and cross-reactivity of clone 52B8 for ILT3 over ILT8. Binding data for the parental mouse anti-human (IgG1) clone 52B8 and Tolerx mouse anti-human ILT3 clone 9B11 are shown with CHO stable cell lines expressing the G223 variant of human ILT3 (A), human ILT8 (B) or rhesus monkey ILT3 (C). The ELISA method is described in the section Materials and Methods.

### ILT3 is an important regulator of dendritic cell activation state and function

We used the antibodies described in Table 1 to interrogate the functional biology of ILT3. At the outset we did not have a well-characterized functional ligand with which to construct a classical blocking assay and therefore we did not know if these particular antibodies would be functionally agonistic, neutral, or antagonistic to ILT3’s presumed inhibitory role in immune cell activation. Our approach then was to develop functional models where ILT3 was implicated as having a role by virtue of up-regulation in that context and on the basis of precedent from published works (21, 42, 43).

As reported by others (42, 43) monocyte-derived dendritic cells (DCs) polarized with IL-10 (Fig. S2A) up-regulated ILT3 surface expression compared to cells not exposed to IL-10 (i.e. immature DCs) and had lower levels of surface CD86 (Fig. S2B). As expected, immature DCs treated with lipopolysaccharide became activated (increased CD86 staining and production of TNFα) but IL-10-treated cells were, in contrast, refractory to LPS (Fig. S2C). Although the effect was modest, we found that addition of anti-ILT3 clone 52B8 during polarization of cells could antagonize the effect of IL-10 with respect to response to LPS for TNFα production (Fig. S2C). Similar experiments were performed using dendritic cells treated with only IL-10 and induction of CD86 expression was observed with addition of anti-ILT3 clone 52B8 (Fig. S2D). This effect of clone 52B8 was consistent across repeat experiments and across donors (data not shown), but it was not unique to clone 52B8. All the four above-mentioned clones could partially antagonize the action of IL-10, although one clone, 51H11, was consistently less active in this assay (Fig. 4A and Table 1). We found that this effect was saturable and dose-dependent in a way broadly consistent with antibody-antigen binding potency across the set of four antibodies (Fig. 4B and Table 1) although a tight correlation between binding potency and functional potency was not evident in this limited set. We were concerned that the activation measured could be an artefact related to labeling of the DCs with an antibody and Fc receptor engagement on a second cell. This did not appear to be the case though because the same activity was observed for human IgG1, effector binding null IgG1(N297A) or IgG4 chimeric variants of clone 52B8 (Fig. 4B). We concluded that the antibodies could be described as antagonists based on these effects on DCs.

**Figure 4.**
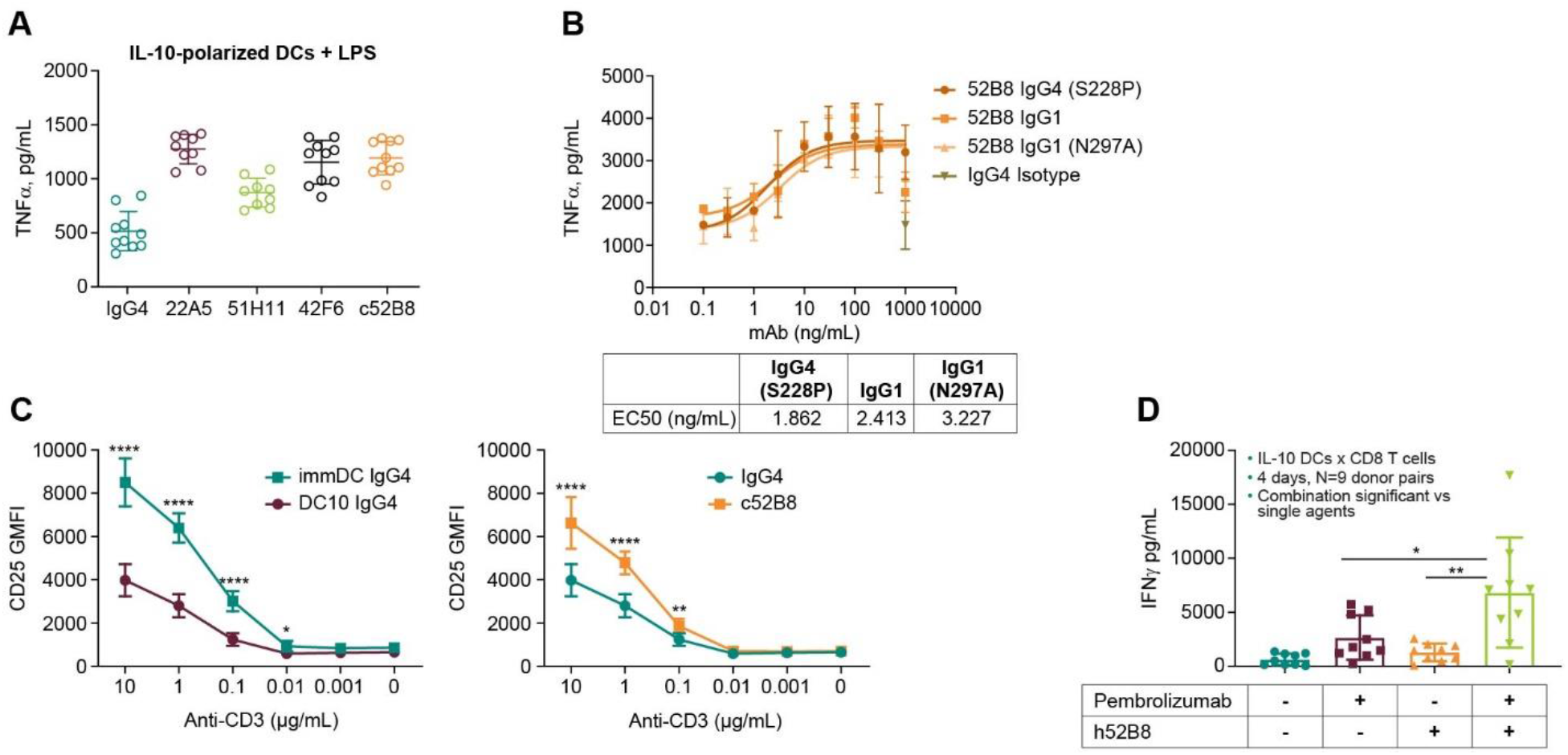
Activation of IL-10-polarized monocyte-derived DCs by anti-ILT3 antibodies. (A) Day 5 Immature DCs were treated with anti-ILT3 antibodies from different epitope bins in the presence of IL-10 & LPS for 42 hours. TNFα concentration in the supernatant was measured.(B) A dose response of chimeric 52B8 with different Fc variants (human IgG4, human IgG1, human IgG1 with N297A mutation) on TNFa production in the DC polarization assay. (C) T cells harvested from mixed lymphocyte reactions with immature or IL-10 DC in the presence of either IgG4 or chimeric 52B8 antibodies were re-stimulated with a titration of anti-CD3 antibody. CD25 expression on CD4^+^ T cells was measured by flow cytometry. Each symbol represents the mean of 3 technical replicates. *: p < 0.05; **: p < 0.01; ****: P< 0.0001, using unpaired t test (D) Combination of 52B8 and pembrolizumab in a mixed lymphocyte reaction using IL-10 polarized DCs & CD8+ T cells, each from three different DC & T cell donors (n=9 pairs) in the presence of indicated antibodies for 4 days where IFNγ was measured in the supernatant. * p val 0.0169, ** p val 0.0011, calculated using one-way ANOVA followed by Tukey’s multiple-comparisons test.

The immediate prediction stemming from these findings is that the DCs exposed to anti-ILT3 antibodies might be better able to prime T cells, this being the rationale for anti-ILT3 antibodies as cancer therapeutics. Indeed, we found early on that clone 52B8 caused an increase in T cell proliferation in a variant of the classical mixed-lymphocyte reaction, but the effect was small, only detectable with many donor pairs (biological replicates) in the experiment and challenging to reproduce with confidence (data not shown). In separate experiments we found that IL-10-polarized DCs had a reduced capacity to prime allogenic CD4+ T cells as revealed in a polyclonal re-stimulation assay format using CD25 surface staining as a marker of T cell activation: T cells primed by IL-10 DCs were less sensitive to anti-CD3 stimulation compared to those primed by immature DCs (Fig. 4C). Notably, the presence of anti-ILT3 antibodies during polarization and priming substantially rescued the priming function when measured in this way (Fig. 4C). Furthermore, in a mixed lymphocyte reaction of IL-10-polarized DCs with allogenic CD8+ T cells, anti-ILT3 treatment during polarization and T cell co-culture could augment the action of PD1 blockade with pembrolizumab on T cell function measured as IFNγ production (Fig. 4D). All these data together indicated a modest but biologically meaningful role for ILT3 in DC function that could be modulated by highly ILT3-specific antibodies.

### ILT3 is important for the suppressive function of cancer cell-educated monocytic myeloid cells

Activation and maturation of dendritic cells rather than tolerance of tumor as self is an important step in immune rejection of a tumor but at the same time non-specific myeloid cell-mediated suppression of T cell function also now appears to be a critical factor in the trajectory of a tumor’s evolution. Like others, we developed an in vitro model to study myeloid-derived suppressor cells. As described in a published paper from the same group (44), we used co-culture with SK-MEL-5 cells to drive acquisition of the suppressor phenotype because we had found that in humanized mice SK-MEL-5 cells can form subcutaneous tumors rich in myeloid cells (Fig. S1C). ILT3 was up-regulated on these cells compared to naïve monocytes (44). MDSCs prepared in this way could potently suppress polyclonal activation of autologous CD8+ T cells but this suppression was abrogated when anti-ILT3 antibody 52B8, either a chimeric IgG4 or a humanized IgG4 variant (Table S8), was included in the co-culture and T cell activation steps (Fig. S3A-B). We have explored this effect in some detail and those data are reported (44). Early in our investigation of ILT3 we conducted a ligand screen under contract by Retrogenix and identified peptidase inhibitor-16 as candidate ligand (data not shown). We had assumed that a relevant ligand interaction would come from PI-16 expression on T cells based on published literature (45). However, inclusion of an antibody to this antigen in our CyTOF panels as a matter of routine resulted in a serendipitous observation that not only could PI-16 be detected on MDSCs but that it was substantively reduced after treatment with anti-ILT3 raising the possibility of a *cis* interaction (Fig. S4A-D). This phenomenon appeared specific because treatment with an anti-ILT4 antibody did not consistently have this effect, despite there being functional ILT4 on these cells (Fig. S4A-D) and also described in a manuscript in review (46). The reduction of PI-16 detection on MDSC after anti-ILT3 treatment was not an interference artefact in staining because PI-16 expression levels on the T cells in the same culture was not changed (Fig. S4E). The effect of clone 52B8 on surface PI-16 levels on MDSCs was dose-dependent and saturable at concentrations consistent with binding and occupancy of ILT3 (Fig. 5A). This finding was consistent across monocyte donors and experiments (Fig. 5B) and was evident following treatment with any of the four anti-ITL3 antibody clones listed in Table 1 (Fig. S4F). Finally, the effect was rapid, it being measurable even with a less than 2-hour incubation with anti-ILT3 antibody at the end of the cancer cell co-culture step (Fig. 5C). These data suggest a proximal complex or signaling effect rather than low surface PI-16 being a feature of the phenotypic changes arising from ILT3 blockade during MDSC differentiation.

**Figure 5.**
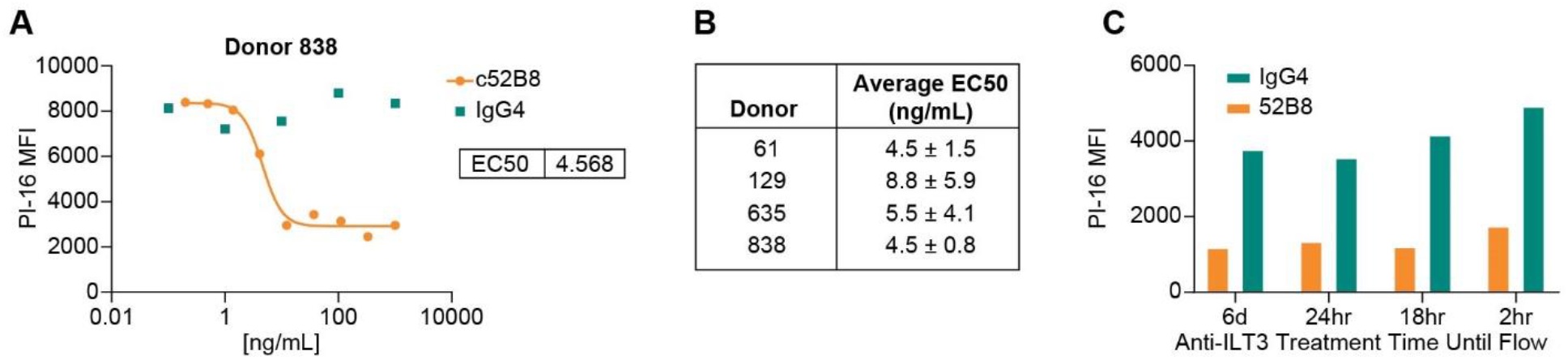
Characterization of anti-ILT3 effect on MDSC phenotype. (A) FACS quantification of PI-16 levels on CD33^+^CD14^+^ MDSCs generated in the presence of varying concentrations of chimeric 52B8 or human IgG4 control. Median fluorescence intensity (MFI) is shown. The data are representative of results from multiple donors and those data are summarized in (B) Average EC50s and SD of n=3 replicates each conducted on separate occasions. (C) Anti-ILT3 antibody 52B8 (1 ug/ml) was added to SK-MEL-5 / monocyte co-cultures at varying times (2, 18, 24 hours and 6 days) before the end of the culture. PI-16 levels on CD33^+^CD14^+^ myeloid cells from the culture were measured by flow cytometry. Data shown are representative of results from 3 donors / co-cultures.

### Anti-ILT3 antibodies slow tumor growth in vivo and cause favorable remodeling of the tumor immune microenvironment

Review of the literature including published data on expression pattern across immune cells in mouse versus human and consideration of sequence and possibly ligand conservation across species led us to conclude that study of mouse GP49b, the nearest homolog of human ILT3, would not be useful. We prioritized *in vitro* human models to assess the role of ILT3 in myeloid cell function and assess utility as a potential therapeutic approach. At the same time, we were curious to investigate the utility of humanized mice bearing human tumors to further our understanding. A variety of approaches have been taken to create and use mice with human immune systems in cancer research (47). We prioritized the hu-NSG model originally developed by Dr. Leonard Shultz from the Jackson Laboratory (48) and used by us previously to study anti-GITR antibodies and anti-ILT4 antibodies (49) (46) (manuscript in review). Initial experiments found that chimeric 52B8 abrogated the growth of SK-MEL-5 tumors (exemplified in Fig. S5A-C) and a subsequent study with humanized 52B8 found the same (Fig. 6A-C). In both studies the effect of anti-ILT3 antibody on in-life tumor volume calculated from caliper measurements was statistically significant. The drug effect was also evident in the weight of the excised tumor mass at necropsy in the humanized study (Fig. 6D).

**Figure 6.**
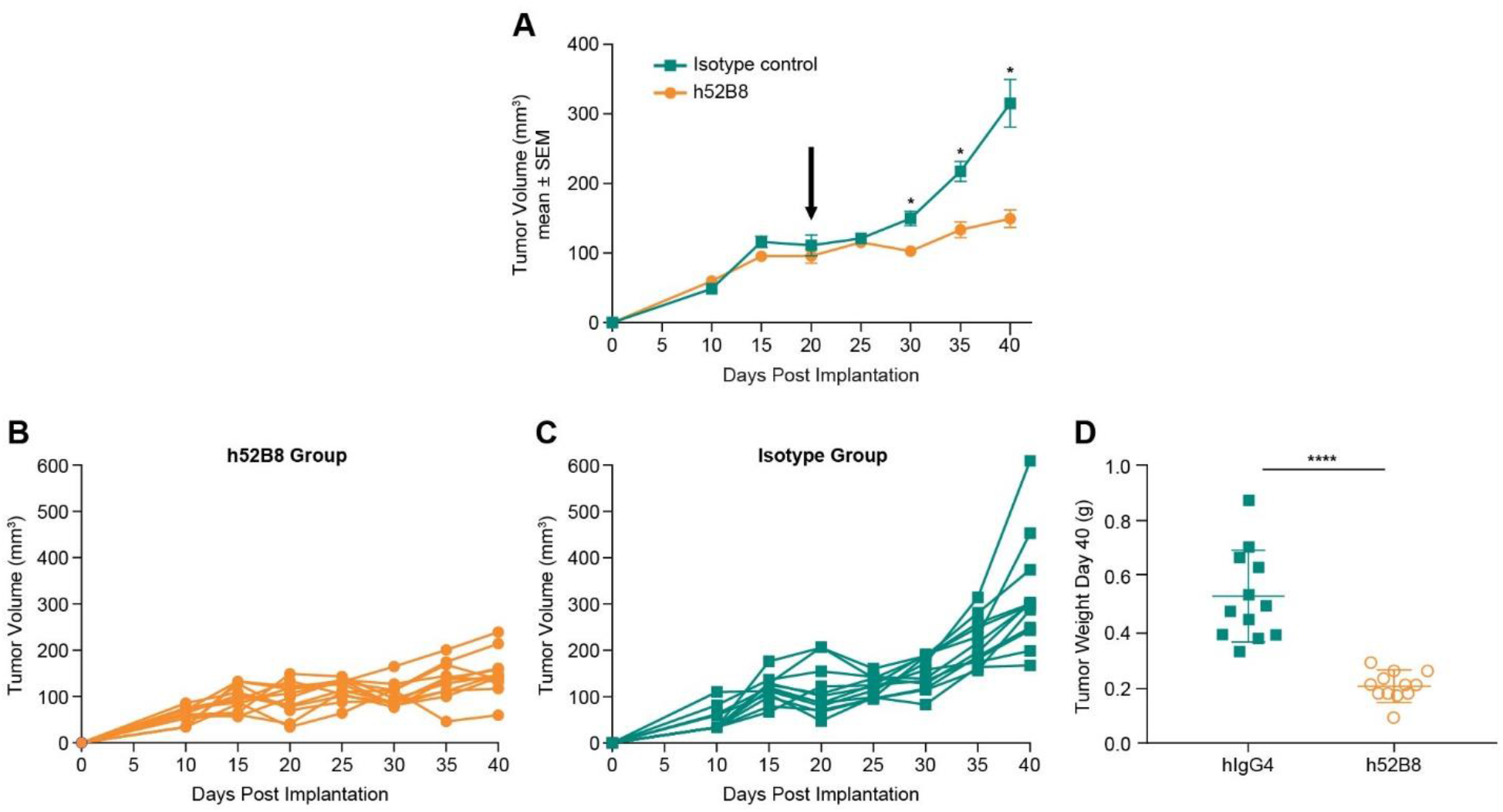
Effect of humanized anti-ILT3 clone 52B8 in humanized mice bearing subcutaneous SK-MEL-5 tumors. Antibodies were dosed at 20 mg/kg once weekly from day 20 (black arrow) and tumor volumes were calculated from caliper measurements at the indicated intervals. **(A)** Mean tumor volumes and SEMs (n=12). Two-sided p-values were estimated from 20,000 random reassignments of animals between the two treatments being compared, with the randomization stratified by donor. To control the familywise error rate across all time points for a given pair of treatments, p-values were multiplicity adjusted by the maxT method of Westfall and Young. A * p-value < 0.05 was used to define statistical significance. **(B and C)** Individual mouse tumor volume data (Spider plots). **(D)** Weights of excised tumors at necropsy, t test used to define significance.

An obvious question was what happened to the tumor immune microenvironment after administration of anti-ILT3 antibodies that caused the slowing of tumor growth. To shed light on this question we conducted new studies to examine the effect of antibody on the myeloid and T cell compartments by CyTOF immune phenotyping (Fig. 7B–E and Tables S6 and S7). The latter studies were carried out in a different vivarium (Boston, MA versus Palo Alto, CA) and it is plainly evident that the size of the effect of antibody on tumor growth is less in these studies (Fig. 7A) than in those in Figs. S5 and Figure 6 and did not reach statistical significance. Nevertheless, we collected the tumors two weeks after treatment for phenotyping. The power of high-dimensional data sets such as this is to look for changes in subsets of cells by unbiased clustering. That is, one does not have to define cell types in advance, but rather the cell types present in the mixture are identified on the basis of over 30 markers using K-means clustering algorithm, allowing sensitivity to biologically meaningful signals that could or would be missed in a classical pre-defined gating approach. We found that *in vivo* anti-ILT3 antibody treatment caused an increase in myeloid cell activation state that matched with our *in vitro* model data (Fig. 7B and C, Table S6). Specifically, we witnessed an increase in two clusters of myeloid cells characterized by concomitant higher expression of HLA-DR, CD86 and CD14 that also express CD11c, possibly activated monocytes or dendritic cells positioned for effective antigen presentation (Fig. 7B and C and Table S6). At the same time, we saw a decrease in clusters of cells characterized by low HLA-DR and CD14 and high immune suppressive markers, such as GITR, PD1, PD-L1 compared to other myeloid cell clusters, which we speculate may be MDSCs, also consistent with our *in vitro* model findings. We also found evidence for remodeling of the tumor immune microenvironment in the T cell compartment (Fig. 7D and E and Table S7). Most notably, we observed with 52B8 treatment increases in cell clusters identified as activated CD4 and CD8 memory and a concomitant decrease in naive Treg cells and antigen experienced T cells with check point inhibitors PD1 and TIGIT. These data are highly complex and certainly open to speculative interpretation, but overall point to an activation of the immune response in 52B8-treated tumors *in vivo* compared to control.

**Figure 7.**
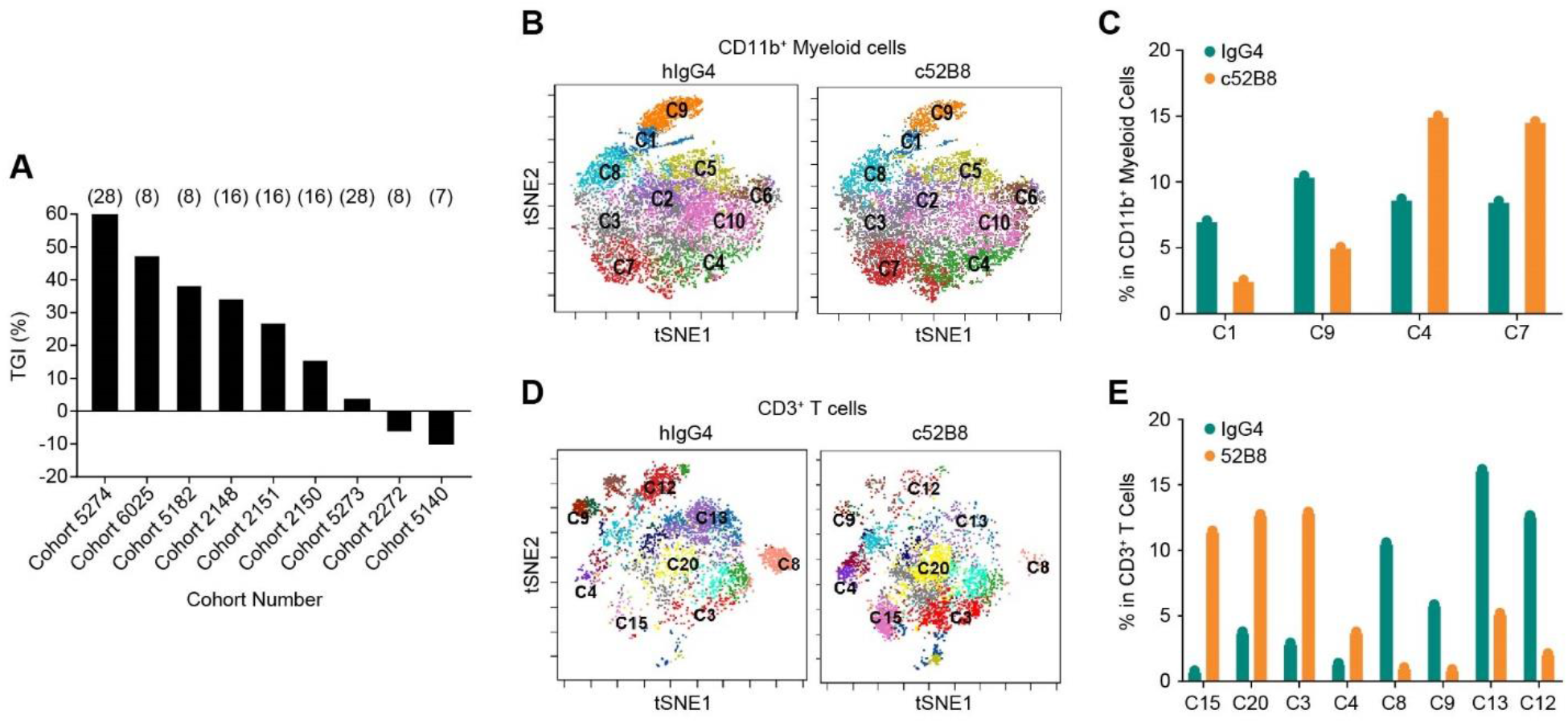
Tumor immune phenotyping before and after treatment with anti-ILT3. **(A)** Compilation of tumor growth inhibition (TGI) in all human cohorts tested using anti-ILT3 chimeric 52B8 (c52B8) in SK-MEL-5 tumor model in the Boston vivarium. Days of antibody treatment are indicated on top of bars **(B)** FlowSOM analysis identified 10 clusters by k-means clustering method for the myeloid cell compartment (n=3 animals concatenated). Distribution of cells across clusters identified in the IgG4- and c52B8-treated groups. **(C)** Numerical representation of data from four clusters with changes detected in (B). **(D)** Cluster assignments by FlowSOM analysis for the T cell compartment (n=3 animals concatenated). Distribution of cells across clusters identified in the IgG4- and 52B8-treated groups. **(E)** Numerical representation of data from eight clusters with changes detected in (D).

## DISCUSSION

In this report we have described initial data that connected ILT3 to cancer immunity and provided an impetus to develop fit-for-purpose pharmacologic tools to investigate the functional role of ILT3 in primary human cell models of tolerance and myeloid immune suppression. Using these tools, namely highly potent and specific antibodies, we found that ILT3 is important in the polarization of monocyte-derived dendritic cells with impact on their ability to prime T cells and we found that ILT3 plays a role in the acquisition of a suppressive phenotype by monocytes when exposed to a cancer cell line. We have corroborated and extended those *in vitro* findings with evidence of remodeling of the tumor immune microenvironment *in vivo*. We were certainly not the first to draw a connection between the ILT family or ILT3 specifically to cancer biology, but our data do provide additional insight and expectations regarding the potential utility of anti-ILT3 antibodies as therapeutic agents. In that sense, our work compliments that of Deng and co-workers that highlighted the likely utility of anti-ILT3 antibodies in the treatment of AML and that of Chen and co-workers that provided a clear rationale for development of anti-ILT4 (LILRB2) antibodies as cancer therapeutics (34, 50). Overall, the model that we subscribe to based on our findings and the accumulated work of others is that ILT3 is an important finger on the scale that controls via inhibitory signaling the balance between immune tolerance and suppression versus recognition of a tumor as not self but rather as an invading pathogen to be targeted and eliminated. In that sense, we hypothesize that in human cancer, therapeutic anti-ILT3 antibodies would tip that balance sufficiently to allow polarization of tumor-resident and infiltrating myeloid cells to support T cell priming and function rather than tolerance of tumor antigens (by APCs) and suppression of T cell function (by MDSCs). Indeed, our in vivo data, while limited at this stage, support exactly that model.

The findings presented in this work leave several questions unanswered and indeed raise additional questions. The first and most important is on signaling – what is ILT3 sensing and how is that signal transducedã Unlike ILT4 and many other immune checkpoint molecules such as PD-1 and TIGIT, there is no definitive and well-characterized canonical ligand identified for ILT3. ApoE has been identified as a candidate ligand (35) (and patent application WO 2016/144728) but we did not detect binding of ApoE to recombinant ILT3 expressed on cells (data not shown). We very much suspect that this relates to either context of the assay or form of the ligand rather than simple non-reproducibility. Similarly, CD166 was reported as a functional ligand for ILT3, albeit with low affinity (36), but again our exploratory binding studies did not recapitulate an interaction. In contrast, we have strong biochemical and cell-binding data for the interaction of ILT4 with HLA-G, and have identified an anti-ILT4 antibody (MK-4830) that effectively blocks this interaction and is currently being tested in the clinic for the treatment of advanced solid tumors (46) (manuscript in review) and (Clinical Trials.gov Identifier: NCT03564691). Other candidate ligands for ILT4, such as β-amyloid, were not substantiated following systematic testing (51-53). Also, we have identified PI-16 and several other undisclosed candidate ligands for ILT3 using a variety of approaches including cell-based membranome screening for *trans* interactors and proximity labeling for *cis*-interactors. In sum, our perspective is that the question of ligand identification for ILT3 is not answered and that it is likely there are multiple ligands relevant in discrete contexts. Intracellular signaling by ILT3 is likely inhibitory given the presence of ITIM motifs in the intracellular domain (32) and several groups have reported that ILT3, like ILT4, can impact canonical tyrosine phosphorylation and cellular activation pathways through SH2 domain containing phosphatases (SHPs) (32-34, 50, 54). We have not investigated intracellular signaling in the primary cell models reported here but this is an important area to study across cell types and states as candidate ligands emerge.

A second key question is the mechanism of action in both the DC setting and the MDSC setting. The default assumption in the DC – MLR setting (Fig. 4) is that the DCs exposed to anti-ILT3 during polarization (and in the MLR culture) are more competent to prime T cells (as witnessed in Fig. 4C-D) and that in either case PD1 blockade facilitates increased T cell activation and the effects are additive. While less interesting than a rigorously synergistic mechanism of action, such an additive effect could nonetheless translate to meaningful improvement in clinical rate and depth of response. We are also conscious of the limitations arising from using monocyte-derived DCs given that these are quite possibly not representative of resident DCs in human tumors. With respect to MDSCs, we have investigated the mechanism of action of ILT3 in some detail and these findings are reported in an accepted manuscript (44). Finally, we have not studied the role of ILT3 in macrophage function. An important next step would be to examine the action of anti-ILT3 antibodies in ex vivo human cancer cultures. The feasibility and utility of this approach was exemplified by Chen and co-workers in their study of ILT4 (50) and could be very valuable in guiding biomarker studies in clinical trials. Tumor efficacy evaluation can be a challenge in humanized mouse models of cancer due to incomplete reconstitution of human immune system and limited cross-talk between human immune cells and mouse parenchyma cells. Nevertheless, we were able to focus our attention in immune activation markers and identify several subsets of activated immune cells unique with anti-ILT3 treatment. It would be very valuable to study anti-ILT3 in additional models, particularly in combination with other therapeutics, notably T cell checkpoint blockers, innate agonists, and chemotherapy.

A third question relates to the importance of soluble ILT3 either as a functional molecule or as a prognostic biomarker in cancer. Suciu-Foca and co-workers first described the increased levels of a soluble form of ILT3 in cancer patients (25), a finding that we have reproduced (Fig. 2B). In that report immunosuppressive activity was attributed to soluble ILT3 in cancer patient serum and other reports described similar activity for ILT3.Fc fusion proteins (29, 30). We were not able to measure any convincing bioactivity for ILT3.Fc fusion proteins and we did not detect any alternative transcripts that could produce a truncated ILT3 either by RT-QPCR or by mining of RNA sequencing data sets. Mass spectrometry studies in cells lines only identified fragments corresponding to the extracellular domain of ILT3. All in all, it is likely that the soluble ILT3 present in serum is simply material shed from myeloid cells en route to elimination. However, we certainly cannot exclude an important and active role for soluble ILT3 in the tumor microenvironment and it may have utility as a patient stratification marker or as a pharmacodynamic biomarker in clinical development.

In summary, it is very clear to us from published work and now with the information presented here that ILT3 plays an important role in the fine balance between tolerance and immune suppression versus immune activation and that the hypothesis that ILT3 blockade will enable productive antitumor immune responses is worth testing in patients suffering from cancers that do not respond well or at all to existing available treatments.

## METHODS

### Analysis of TCGA and pembrolizumab response data

The Cancer Genome Atlas (TCGA) database was used for analysis of clinical relevance. RNA-sequencing data for 9963 tumors and somatic alterations data for 6384 tumors were obtained through TCGA portal (https://portal.gdc.cancer.gov/) as of September 2015 (The Cancer Genome Atlas, TCGA Data Portal; https://tcga-data.nci.nih.gov/docs/publications/tcga/). The expression data were log10 transformed. Spearman correlation was used to determine the correlation and Wilcoxon rank-sum test was used to calculate P value. Statistical analyses and visualizations were performed with Matlab R2010b Version 7.11.2.

TMB cutoff for the pan-tumor clinical cohort were the Youden Index value derived in AUROC analysis. An additional, exploratory, pan-tumor TMB threshold was derived by using TMB and GEP data, similar to a previously described method (A. Panda, et al Identifying a clinically applicable mutational burden threshold as a potential biomarker of response to immune checkpoint therapy in solid tumors. JCO Precis. Oncol. 2017, 10.1200/PO.17.00146 (2017). pmid:29951597 (ref 20 in science paper)).

### Measurement of surface expression of ILT3 in human tumors

Single cell suspensions of tumors were obtained by fine mincing with a scalpel, followed by a 30-minute incubation at 37°C in digestion medium containing 8 mL of Roswell Park Memorial Institute 1640 medium (RPMI 1640), 40 µL of 100 mg/mL collagenase I, and 320 µL of 10,000 U/mL DNase I. Samples were filtered, red blood cells were removed by ACK lysis, and cell numbers were counted using either a Hemocytometer or the ViCell Cell Viability Analyzer (Beckman Coulter).

One million (10^6^) viable cells per tube were stained for viability using Fixable Viability Dye eFluor 506 in DPBS followed by fluorescently-labeled mAb against CD11b, CD3, CD66b, CD33, CD45, CD56, CD14, anti-ILT3 clone ZM4.1, or mouse IgG1 isotope control as listed below in FACS staining buffer on ice. All samples were acquired on a LSR II flow cytometer using FACSDiva™ software (BD Biosciences). All flow cytometry data was analyzed using FlowJo version 10.0.7 or above (FlowJo, LLC). Expression of ILT3 was calculated as the frequency of cells showing greater fluorescence signal when incubated with anti-ILT3 clone ZM4.1 than those incubated with the mouse IgG1 isotype control mAb.

Antibodies used – CD11b (BUV395 BD Biosciences, #56553, clone M1/70, 1.25 uL, human/mouse), CD3 (BUV737 BD Biosciences, #564307, clone UCHT1, 5 uL, human), CD66b (FITC, BioLegend, #305104, clone G10F5, 5 uL, human), CD33 (PE/CF594, BD Biosciences, #562492, clone WM53, 5 uL human), CD45 (AF700, Biolegend, #304024, clone HI30, 2 uL, human), CD56 (BV605, BD Biosciences, #562780, clone NCAM16.2, 5 uL, human), CD14 (BV650, BioLegend, #301836, clone M5E2, 5 uL, human), ILT3 (PE/Cy7, BioLegend, #333012, clone ZM4.1, 5 uL, human), Mouse IgG1 Isotype Control (PE/Cy7, BioLegend, #400126, clone MOPC-21, 2.5 uL, NA)

### CyTOF immune-phenotyping

Resected tissue from human mesothelioma and lung adenocarcinoma patients were provided to Merck & Co., Inc., Kenilworth, NJ, USA through a collaboration with Dr. Raphael Bueno at Brigham and Women’s Hospital, with IRB approval. For the mesothelioma sample, tissue was diced into smaller pieces and digested into a single cell suspension using the protocol from Miltenyi Tumor Digestion Kit, (P# 130-095-929). For the lung adenocarcinoma sample, tissue was diced into small pieces and incubated at 37C for 30 minutes with intermittent shaking in 0.5 mL digestion medium made by adding 8 mL of DMEM + 40 uL of 250 U/mL Collagenase Type 1+ 320 uL of 10,000 U/mL DNAse I. Tissue particles were disrupted using a syringe plunger and cells were filtered through a 100 μm strainer over a 50 mL conical tube.

SK-MEL-5 tumors in hu-NSG mice were collected 29 days after tumor cell implantation (7 days after the second dose of antibody treatment). Single cell suspension was generated using the Miltenyi Tumor Digestion Kit (P# 130-095-929).

Single tumor cell suspensions were stained using a panel of CyTOF antibodies (see supplemental tables), profiled on the Helios mass cytometer, and analyzed using Cytobank.

Cytof Panel used in SK-MEL-5 hu-NSG model: Antigen (Metal-tag, Clone, Vendor, Catalog#)

CD3 (154Sm, UCHT1, Fluidigm, #3154003B), CD4 (145Nd, RPAT4, Fluidigm, #3145001B), CD8 (146Nd, RPAT8, Fluidigm, #3146001B), CD11b (209Bi, ICRF44, Fluidigm,#3209003B), CD11c (147Sm, Bu15, Fluidigm, #3147008B), CD14 (148Nd, RMO52, Fluidigm, #3148010B), CD15 (164Dy,W6D3, Fluidigm, #3164001B), CD19 (142Nd, HIB19, Fluidigm, #3142001B), CD33 (169Tm, WM53, Fluidigm, #3169010B), CD34 (166Er,581, Fluidigm, #3166012B), CD45 (89Y, HI30, Fluidigm, #3089003B), CD45RO (165Ho, UCHL1, Fluidigm, #3165011B) CD45RA (143Nd, HI100, Fluidigm, #3143006B), CD66b (152Sm, 80H3, Fluidigm, #3152011B), CD103 (172Yb, BerACT8, Fluidigm, #3151011B), CD25 (149Sm, 2A3, Fluidigm, #3149010B), CD27 (155Gd, L128, Fluidigm, #3155001B), CD28 (160Gd, CD28.2, Fluidigm, #3160003B), CD69 (162Dy, FN50, Fluidigm, #3162001B), CD86 (150Nd, IT2.2, Fluidigm, #3150020B), CD127 (176Yb, A019D5, Fluidigm, #3176004B), CD137 (158Gd, 5F4, Fluidigm, #3158022B), CD152 (170Er, 14D3, Fluidigm, #3170005B), CD154 (168Er, 2431, Fluidigm,#3168006B), CD197(CCR7) (167Er, G043H7, Fluidigm, #3167009A), CD226 (171Yb, DX11, Fluidigm, #3171013B), CD278(ICOS) (151Eu, C398.4A, Fluidigm, #3151020B) HLA-DR (174Yb, L243, Fluidigm, #3174001B), CD274(PD-L1) (156Gd, 29E.2A3, Fluidigm, #3156026B), CD279(PD-1) (175Lu, EH12.2H7, Fluidigm, #3175008B), TIGIT (153Eu, MBSA43, Fluidigm, #3153019B), GITR(CD357) (159Tb, 621, Fluidigm, #3159020B), ILT3(CD85k) (173Yb, ZM4.1, eBioscience,#16-5139-85), ILT4 (161Dy, 42D1, Fluidigm, #3161019B), HLA-G (163Dy, MEM-G/9, Fluidigm, #ab7758), HLA-ABC (144Nd, W6/32, Fluidigm, #3144017B), HLA-A2 (141Pr, BB7.2, Thermo Fisher, MA1-80117)

Cytof Panel used for human tumor study: Antigen (Metal-tag, Clone, Vendor, Catalog#)

CD3 (170Er, UCHT1, Fluidigm, #3170001B), CD4 (145Nd, RPAT4, Fluidigm, #3145001B), CD8 (162Dy, RPAT8, Fluidigm, #3162015B), CD11b (209Bi, ICRF44, Fluidigm,#3209003B), CD11c (147Sm, Bu15, Fluidigm, #3147008B), CD14 (148Nd, RMO52, Fluidigm, #3148010B), CD19 (142Nd, HIB19, Fluidigm, #3142001B), CD25 (169Tm, 2A3, Fluidigm, # 3169003B), CD27 (155Gd, L128, Fluidigm, #3155001B), CD28 (160Gd, CD28.2, Fluidigm, #3160003B), CD33 (158Gd, WM53, Fluidigm, #3158001B), CD38 (167Er, HIT2, Fluigim, #3167001B), CD40 (165Ho, 5C3, Fluidigm, #3165005B), CD45 (89Y, HI30, Fluidigm, #3089003B), CD45RO (164Dy, UCHL1, Fluidigm, #3164007B), CD45RA (153Eu, HI100, Fluidigm, #3153001B), CD56 (176Yb, NCAM16.2, Fluidigm, #3176008B), CD64 (146Nd, 10.1, Fluidigm, #3146006B), CD66b (152Sm, 80H3, Fluidigm, #3152011B), CD68 (171Yb, Y1/82A, Fluidigm, #3171011B), CD69 (144Nd, FN50, Fluidigm, #3144018B), CD86 (150Nd, IT2.2, Fluidigm, #3150020B), CD123 (151Eu, 6H6, Fluidigm, #3151001B), CD127 (149Sm, A019D5, Fluidigm, #3149011B), CD141 (166Er, M80, Fluidigm, #3166017B), CD163 (154Sm, GHI/61, Fluidigm, #3154007B), CD172a/b (163Dy, SE5A5, Fluidigm, #3163017B), CXCR3 (156Gd, G025H7, Fluidigm, #3156004B), CD278(ICOS) (143Nd, C398.4A, Fluidigm, #3143025B), HLA-DR (174Yb, L243, Fluidigm, #3174001B), PI-16 (141Pr, NA, Merck & Co., Inc., Kenilworth, NJ, USA, NA), CD274(PD-L1) (159Tb, 29E.2A3, Fluidigm, #3159029B), CD279(PD-1) (175Lu, EH12.2H7, Fluidigm, #3175008B), CD273(PDL-2) (172Yb, 24F.10C12, Fluidigm, #3172014B), ILT3(CD85k) (112Cd, ZM4.1, eBioscience, #16-5139-85), ILT4 (161Dy, 42D1, Fluidigm, #3161019B), Ki67 (168Er, Ki-67, Fluidigm, #3168001B), Vd1 (110Cd, TS8.2, Invitrogen, TCR1730), Vd2 (111Cd, B6, Biolegend, #331402)

### Measurement of soluble ILT3 in human plasma

Human plasma samples were acquired from Bioreclamation or Sanguine. Levels of sILT3 in serum samples from healthy donors and cancer patients were measured using an electrochemiluminescence (ECL) assay on the Meso Scale Discovery (MSD) platform. The lower limit of quantitation (LLOQ) of the assay is 0.002 ng/mL. The capture reagent was immobilized biotinylated mouse anti-ILT3 clone ZM4.1 (ebioscience). The detection reagent was Sulfo-labelled-mouse anti-human ILT3 (VEGF leader]-LILRB4_H/ILT3_H-[His9] AA#22-259, Lot #41AIT). The signal was read on the SECTOR Imager 6000 (Meso Scale Discovery).

### Selectivity and cross-reactivity of clone 52B8 for ILT3 over ILT8

Mouse anti-human ILT3 antibodies were tested for binding to human ILT3, and cross-reactivity to Rhesus monkey ILT3, human ILT5, human ILT7, human ILT8, and human ILT11 expressing CHO-K1 cells using a cell - based ELISA format. CHO-K1 cells were plated in 96-well tissue culture plates in 50 uL of DMEM / F12, 10 % BCS and gentamycin (CHO-K1 media). Cells were plated at either 2×10^4^ cells/well two days prior to the assay or 4×10^4^ cells/well one day prior to the assay. Media was removed from the wells prior to adding the test samples. Purified antibody was serially-diluted in CHO-K1 media and added to the CHO - K1 plates. The samples were incubated at room temperature for 30 - 60 minutes and plates were washed three times with PBS / 0.5 % Tween-20 using the cell wash program on the Biotek EL405x Select CW plate washer. Binding was detected using an HRP-conjugated goat anti-mouse IgG (Southern Biotech, cat # 1031-05) secondary antibody added at a 1: 2000 dilution in CHO-K1 media and incubated at room temperature for 30 - 60 minutes. Assay plates were washed and developed with TMB and stopped with TMB stop solution (KPL, cat# 50-85-06). The absorbance at 450 nm - 620 nm was determined. Mouse IgG1 (MIgG1) served as a control.

### Monocyte Derived Dendritic Cells Generation, Polarization, and Phenotyping

CD14+ monocytes were selected from frozen human PBMC using anti-CD14 microbeads according to manufacturer’s instructions (Stemcell Technologies). The isolated monocytes were counted and resuspended in growth medium (Stemcell Technologies, DC differentiation Medium, serum-free) supplemented with GM-CSF (1000 U/mL) and IL-4 (1000 U/mL) or GM-CSF & IL-4 (20 and 40 ng/mL, R&D Systems) and seeded into 24-well low binding plates at 1 × 10^6^ cells/well. Cells were incubated for 5 days at 37°C (5% carbon dioxide) and non-adherent cells were collected as monocyte-derived dendritic cells.

Day 5 immature DCs were then further cultured for 42 hours with addition of IL-10 (50 ng/mL) and/or LPS (1 ug/mL) with or without anti-ILT3 mAbs. TNFα in the culture supernatant was measured on the MSD platform (Mesoscale #K15049D). Cells were collected and surface expression of CD86 (Biolegend, #305420), ILT3 (Biolegend, #333016), HLA-DR (Biolegend, #307620) or PD-L1 (Biolegend, #329714) was measured by flow cytometry.

### IL-10 Polarized Dendritic Cell and Lymphocyte Coculture

CD3+ T cells and IL10-tolerized or untreated dendritic cells from allogeneic healthy human PBMCs were co-cultured and incubated with either 10 µg/mL human IgG4 (hIgG4) or 10 µg/mL 52B8. Following 7 days, CD3+ T cells were enriched using magnetic beads and re-stimulated with anti-CD3 mAbs (eBioscience, clone OKT3) for 3 days. CD25 levels on CD4 was assessed by flow cytometry (Biolegend, clone BC96, #302612).

IL-10 polarized human DCs (3×10^4/well) were incubated with allogeneic CD8+ T cells (1.5 × 10^5/well) in T cell media (Stem Cell Technologies) in 96 well flat bottom tissue culture plates for 4 days. CD8+ T cells were isolated from freshly-thawed PBMCs using the human CD8+ T cell isolation kit (Stem Cell Technologies). Anti-ILT3 52B8 (1µg/mL) and/or pembrolizumab (2µg/mL) were added in the culture. Human IgG4 Ab was used as isotype controls. IFNγ levels in the culture supernatant were determined by MesoScale Discovery kits (K151QO Series) using MesoScale Discovery SECTOR 6000. Curve fitting was performed within Mesoscale Discovery Workbench 4.0 software using Four Parameter Logistic fit to calculate concentrations of IFNγ.

### In vitro activity of anti-ILT3 in myeloid-derived suppressor cells and T cell co-culture assay

Healthy human PBMCs from individual donors (1 ×10^6 cells/mL) were cultured in complete cell culture medium (RPMI-1640 with 10% heat-inactivated fetal bovine serum [Hyclone, Inc.], 2 mM L-glutamine, 50 units/mL each of penicillin/streptomycin) in culture flasks with SK-MEL-5 tumor cells in the presence of 20 ng/mL GM-CSF at 37°C for 7 days. SK-MEL-5 tumor cells were seeded to achieve confluence by day 7 (approximately 1:50 ratio with PBMC). Co-cultures were incubated with 1 µg/mL hIgG4 or 1 µg/mL anti-ILT3 antibody (c52B8 or h52B8). Media supplemented with fresh GM-CSF and hIgG4 or anti-ILT3 was changed on day 4. After the 7-day co-culture period, the cells were harvested using Detachin solution, and CD33+ myeloid cells were isolated using anti-CD33 magnetic microbeads and LS column separation (Miltenyi Biotech) according to the manufacturer’s instructions.

To assess the single agent activity of anti-ILT3 on reversal of T cell suppression induced by MDSCs, purified autologous CD8+T cells (1 × 10^5 cells per well) were cultured alone or co-cultured with CD33+ myeloid suppressive cells at the ratio of 4:1 (T cell:MDSC) in 96-well U-bottom plates for 30 minutes, in the presence of 1 µg/mL hIgG4 or 1 µg/mL anti-ILT3 antibody (c52B8 or h52B8). After the pre-incubation, T cell proliferation was induced with anti-CD3/CD28 beads (Thermo Scientific #11161D) and 100 U/mL IL2 (Thermo Scientific cat# PHC0026) and incubated at 37°C for 3 days. IFN-γ levels in culture supernatants were determined using Mesoscale Discovery kits according to the manufacturer’s protocol.

### Modulation of PI-16 binding by anti-ILT3 in myeloid-derived suppressor cells

Human PBMCs were cultured at a 50:1 concentration with SK-MEL-5 tumor cells in complete culture medium containing GM-CSF (20 ng/mL). In some experiment cells were treated for 7 days with varying concentrations of different anti-ILT3 antibodies, anti-ILT4 clone 1E1, or human IgG4 control in a 96-well format at 37°C (5% carbon dioxide). In other experiments, chimeric 52B8 and human IgG4 control concentration was fixed (1 ug/mL) and duration of antibody treatment during PBMC tumor education varied (ranging from 6 days to 2 hours prior to cell collection). Following the incubation period, all cells were collected and stained for flow cytometry. To address the possibility of staining interference, anti-ILT3 52B8 or human IgG4 control was added to untreated collected cells on ice for 10 minutes prior to staining. Cells were stained with fixable viability dye BV510 (BD Biosciences, Catalog # 564406) before antibody staining, where CD45 APC-H7, (BD Biosciences #560178), CD14 BV785 (Biolegend #301839), CD33 PE (Biolegend #303404), ILT3 PerCP-Cy5.5 (Biolegend #333014), CD3 BV421 (Biolegend #300434), and a Merck & Co., Inc., Kenilworth, NJ, USA PI-16 antibody labeled with AF647 (Invitrogen #A20186) were added to FACs staining buffer on ice. Samples were collected on a BD Fortessa Cytometer and analyzed in FCS Express 7 Software. Surface PI-16 MFI or percent positive cells was measured from CD33+ CD14+ gated MDSCs. For experiments where dose responses were generated the data was plotted in graphpad prism using a log vs response variable slope (four parameters).

### SK-MEL-5 tumor in hu-NSG model

Immunodeficient NSG mice were reconstituted with human hematopoietic stem cells (CD34^+^) from 2 – 3 different cord blood donors at The Jackson Laboratory (Bar Harbor, ME). After mice were confirmed to harbor peripheral human CD45^+^ immune cells (>25% of total blood leukocytes), they were shipped to either Palo Alto or Boston site where they were subcutaneously (SC) inoculated with the human skin melanoma derived tumor line SK-MEL-5 (1×10^6^ cells/mouse) in their flanks at approximately 20 weeks of age. Treatment started when mean tumor size was approximately 100 mm^3^. Mice were randomized into groups. Tumor-bearing mice were injected SC or IP with 20 mg/kg of chimeric or humanized anti-ILT3 clone 52B8 or a hIgG4 isotype control mAb every 5 or 7 days. Tumors were measured using a caliper and tumor volume was calculated as length x width x ½ width. Tumor growth inhibition (TGI) was calculated as % of (isotype treated tumor volume increase – 52B8 treated tumor volume increase)/isotype treated tumor volume increase.

### Statistics

Data were compared using a one-way ANOVA followed by Tukey’s multiple-comparisons test or one-way ANOVA followed by Dunnett’s correction when all data were compared with a control group. One sided or two tailed paired or unpaired t test was used. For the in vivo studies two-sided p-values were estimated from 20,000 random reassignments of animals between the two treatments being compared, with the randomization stratified by donor. To control the familywise error rate across all time points for a given pair of treatments, p-values were multiplicity adjusted by the maxT method of Westfall and Young. Data was considered statistically significant with p value < 0.05. Analysis was performed in Graph Pad Prism 8 software or using R, version 3.5.1.

### Study Approval

All animal work was reviewed and approved by IACUC at either Palo Alto or Boston site (Merck & Co., Inc., Kenilworth, NJ, USA) before experiments were conducted. Use of human tumor tissue samples for profiling were used with IRB approval.

## AUTHOR CONTRIBUTIONS

P.E.B wrote the manuscript with support from J.Z.H. and A.P.

A.P., G.A., L.B., J.A., A.B., M.C., C.C., Y.C., H.C., D.C., X.D., L.F.D., B.H., H.I., B.J.S., V.J., C.M., Y.Q., L.S., P.S., K.V., D.C.W., C.Z designed, conducted and analyzed experimental studies.

A.L, J.L performed informatics analysis on TCGA and pembrolizumab response datasets

R.B, R.D.L, P.J.T, K.V. contributed provided human tumor tissue.

P.E.B, M.M, J.Z.H., M.R. designed and directed the project

