## Supplemental Figures & Tables for "Antibodies to ILT3 abrogate myeloid immunosuppression and enable tumor killing"

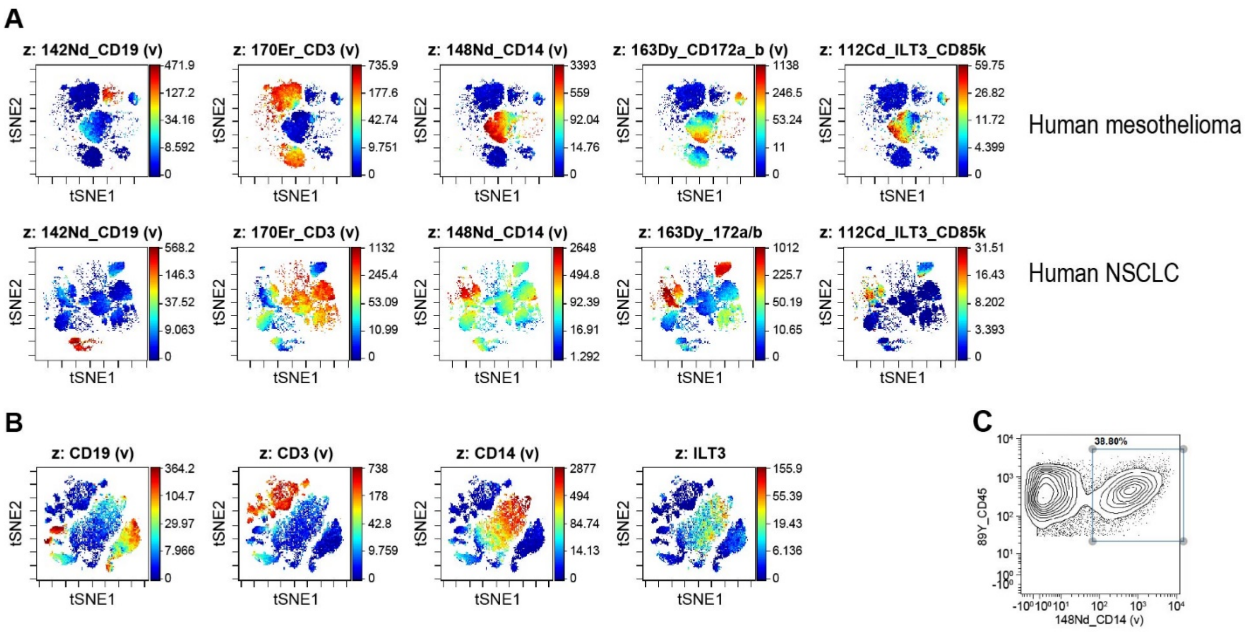


**Figure S1. Expression of cell surface ILT3 protein in tumors measured using CyTOF. (A)** Human mesothelioma and NSCLC fresh biopsy, one sample representative of samples from two patients. **(B)** SK-MEL-5 subcutaneous tumors grown in humanized mice, concatenated viSNE cluster analysis of three CyTOF FCS files from individual mice euthanized at day 29 after implantation. In both cases, the total human CD45+ cell population is included in the viSNE plots, and the z-axis color scale corresponds to signal intensity for the indicated antigen (CD19, CD3, CD14 or ILT3). **(C)** Percentage of CD14+ myeloid cells in total human CD45+ TILs in SK-MEL-5 tumors.

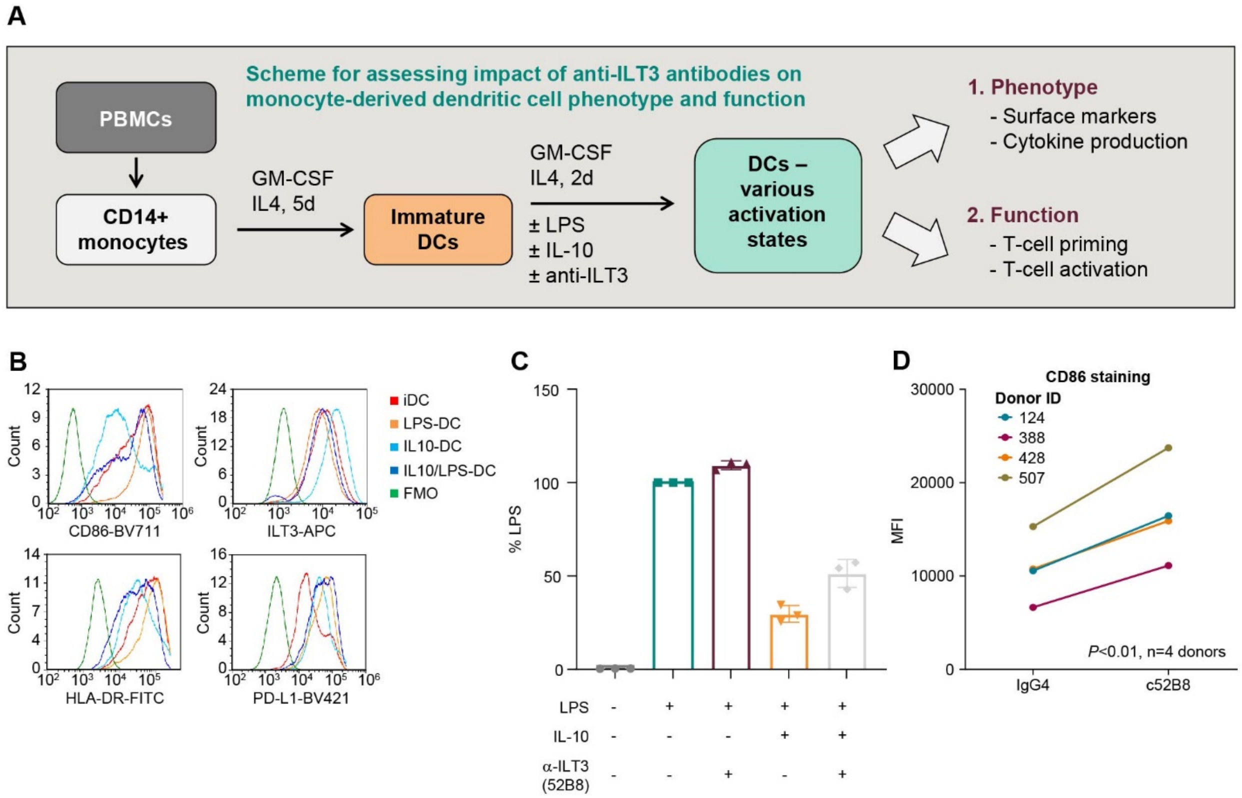


**Figure S2. Phenotype of human monocyte-derived DCs polarized with IL-10.** (A) Schematic for derivation and characterization of monocyte derived dendritic cells. CD14+ monocytes were enriched from PBMCs and cultured in GM-CSF & IL4 for 5 days. Cells are either left untouched (Immature DCs) or cultured for 2 additional days in the presence of LPS, IL-10, LPS + IL-10 and/or anti-ILT3 antibody. Cells are phenotyped for expression of surface markers and cytokine production or used in functional studies. (B) Surface expression of CD86, ILT3, HLA-DR or PD-L1 on DCs treated under noted conditions. IL-10 DCs showed increased expression of ILT3 and lower levels of CD86 relative to immature DCs. (C) Effect of anti-ILT3 on TNF alpha production or (D) CD86 expression levels in DCs. Significance was determined using a one sided paired t test.

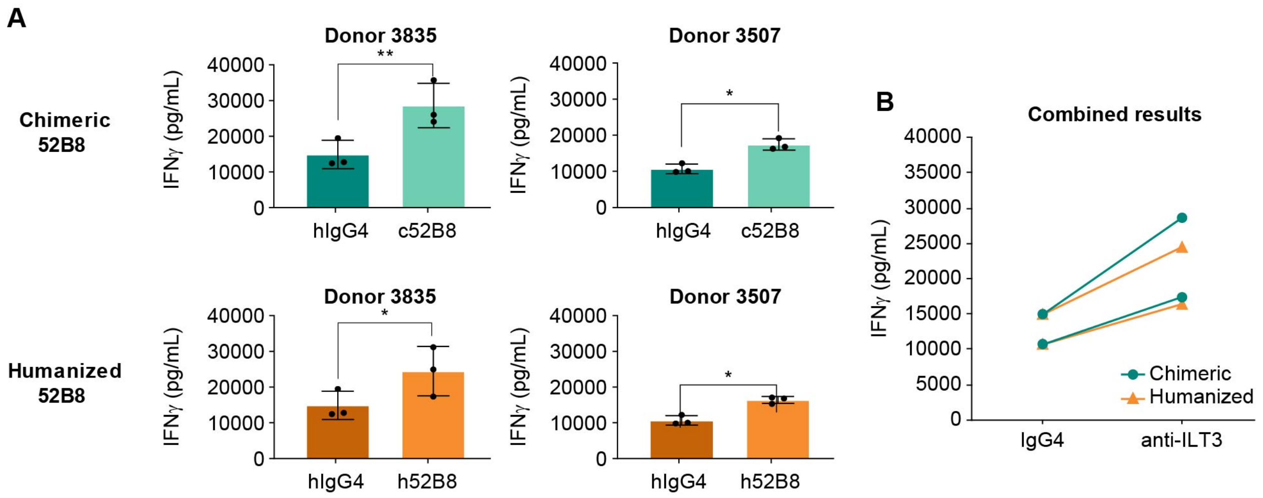


**Figure S3. Characterization of anti-ILT3 effect on MDSC function.** (A) IFNg levels in 3-day co-cultures of MDSC and autologous CD8+ T cell activated with anti-CD3/CD28 beads and IL2. Anti-ILT3 antibody (c52B8 or h52B8) and isotype control antibody (1 µg/mL) were used in both the 7-day culture that MDSC were generated and the 3-day co-culture of MDSC with CD8+ T cells (T cell to MDSC ratio of 4:1). The results are expressed as average ± standard deviation from 3 technical replicates for each donor, and P value was calculated using paired t test (one-tailed), *p <0.05, **p <0.01. (B) Combined data set from (A). The effect of anti-ILT3 antibody (chimeric or humanized) treatment was significant (p<0.01, two tailed paired t-test).

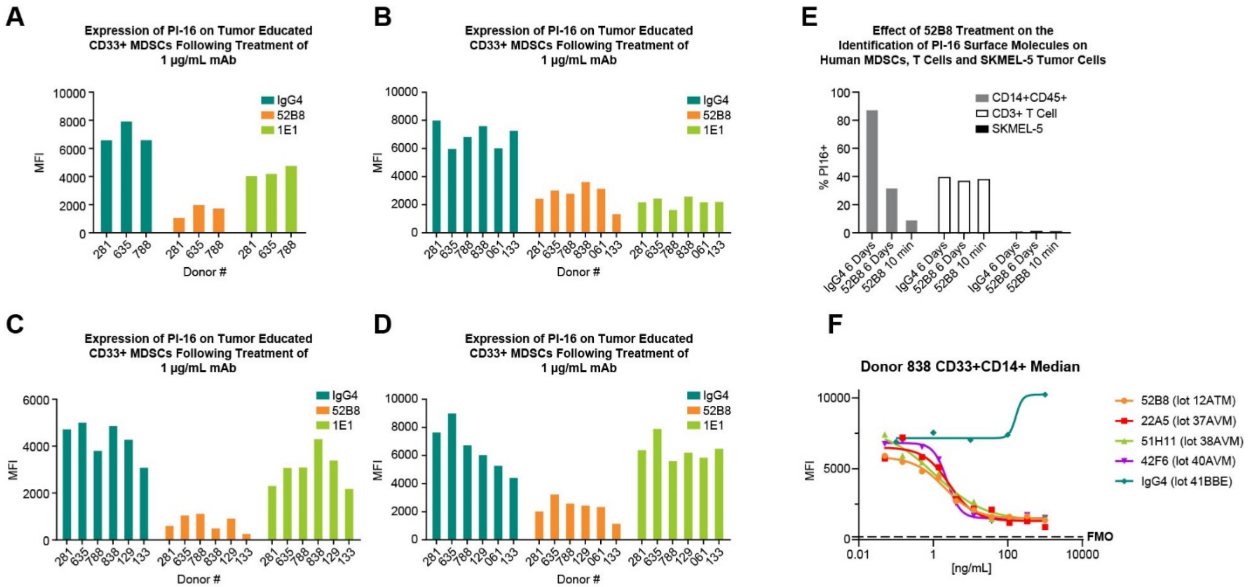


| | 52B8 (lot 12ATM) | 22A5 (lot 37AVM) | 51H11 (lot 38AVM) | 42F6 (lot 40AVM) |
|---|---|---|---|---|
| EC50 | 2.011 | 2.688 | 1.105 | 2.339 |

**Figure S4. Effect of anti-ILT antibodies on the surface expression of PI-16 on MDSCs. (A – D)** Four separate experiments where monocytes from multiple donors were co-cultured with anti-ILT antibodies or human IgG4 control for 7 days and PI-16 was measured on CD33+ cells by flow cytometry. ILT3 – clone 52B8, ILT4 – clone 1E1, 1 ug/ml each. Clone 1E1 was also tested at 10 ug/ml in some donors with no difference in effect (data not shown). **(E)** Control data to address the possibility of inhibition of detection of PI-16 by 52B8. **(F)** Monocytes were co-cultured with SKMEL5 cells for 7 days in the presence varying concentrations of the four different anti-ILT3 antibodies described in Table 1 and PI-16 was measured on CD33+ cells.

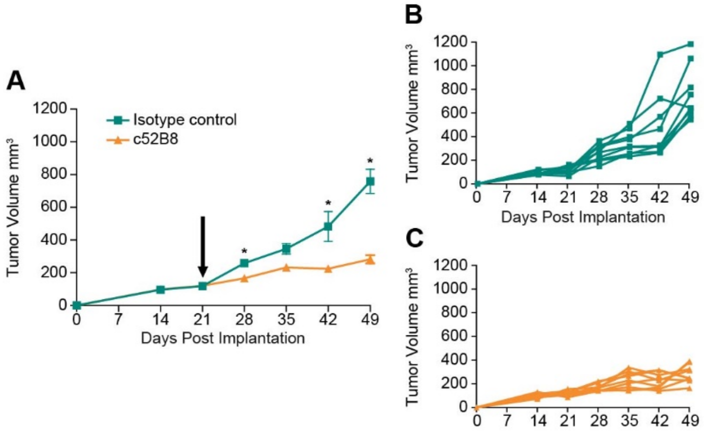


**Figure S5. Effect of anti-ILT3 clone 52B8, chimeric human IgG4 in humanized mice bearing subcutaneous SK-MEL-5 tumors.** Antibodies were dosed at 20 mg/kg once weekly from day 21 (black arrow) and tumor volumes were calculated from caliper measurements at the indicated intervals. **(A)** Mean tumor volumes (n=9) and SEMs. **(B and C)** Individual mouse tumor volume data (Spider plots). Two-sided p-values were estimated from 20,000 random reassignments of animals between the two treatments being compared, with the randomization stratified by donor. To control the familywise error rate across all time points for a given pair of treatments, p-values were multiplicity adjusted by the maxT method of Westfall and Young. A * p-value of less than 0.05 was used to define statistical significance.

**Table S6. Phenotypic assignment and CyTOF marker intensity of discrete myeloid cell clusters in SK-MEL-5 tumors from Figure 7 B & C.**

| Cluster # | Cell Type | Specific Markers | CD11b | CD11c | CD14 | CD33 | CD66b | CD15 | CD86 | ICOS | CD103 | GITR | HLADR | PDL1 | PD1 | ILT3 |
|---|---|---|---|---|---|---|---|---|---|---|---|---|---|---|---|---|
| 1 | Monocytic MDSC | CD14$^{low}$CD33$^{low}$HLA-DR$^{low}$CD86$^{-}$ | 14.59 | 11.75 | 5.22 | 6.84 | 0.27 | 1.12 | 4.54 | 1.11 | 2.1 | 7.99 | 42.52 | 5.82 | 3.28 | 0.46 |
| 9 | Lymphoid-derived Dendritic cells | CD14$^{low}$CD33$^{+}$ICOS$^{+}$CD103$^{+}$GITR$^{+}$PD-L1$^{+}$PD-1$^{+}$HLA-DR$^{low}$ | 20.08 | 24.21 | 15.59 | 29.9 | 1.55 | 4.17 | 12.16 | 9.52 | 14.67 | 21.51 | 106.14 | 27.95 | 18.22 | 2.09 |
| 4 | Monocytes | CD11c$^{+}$CD14$^{high}$CD33$^{+}$CD15$^{+}$CD86$^{+}$HLA-DR$^{high}$ | 103.45 | 210.91 | 1034.66 | 111.73 | 2.35 | 24.28 | 23.48 | 1.7 | 2.67 | 2.83 | 1151.61 | 5.69 | 5.07 | 22.61 |
| 7 | Monocytes | CD11c$^{high}$CD14$^{high}$CD33$^{+}$CD15$^{+}$CD86$^{+}$HLA-DR$^{high}$ | 67.43 | 418.93 | 1194.21 | 74.09 | 3.28 | 28.85 | 28.55 | 2.19 | 2.86 | 3.67 | 1147.27 | 7.24 | 4.96 | 15.66 |

**Table S7. Phenotypic assignment and CyTOF marker intensity of discrete T cell clusters in SK-MEL-5 tumors from Figure 7 D & E.**

| Cluster # | Cell Type | Specific Markers (cell lineage, activation, checkpoint) | CD4 | CD8 | CD127 | CD25 | CD45RA | CD45RO | CD69 | CCR7 | HLA-DR | PD1 | TIGIT |
|---|---|---|---|---|---|---|---|---|---|---|---|---|---|
| 15 | Activated $CD4^+$ memory T cells | $CD4^+$ $CD45RO^+$ $CD69^+$ $HLA\text{-}DR^+$ | 99.06 | 8.49 | 3.91 | 2.57 | 0.98 | 202.95 | 52.89 | 1.94 | 61.92 | 50.35 | 5.98 |
| 20 | Activated $CD4^+$ memory T cells | $CD4^+$ $CD45RO^+$ $CD69^+$ $HLA\text{-}DR^-$ | 93.84 | 7.61 | 5.97 | 1.86 | 1.9 | 103.96 | 17.1 | 16.13 | 7.24 | 31.96 | 4.33 |
| 3 | "effector" $CD4^+$ Treg cells | $CD4^+$ $CD127^-$ $CD25^{intermediate}$ $CD45RO^+$ ICOS+ $HLA\text{-}DR^{high}$ | 126.64 | 10.2 | 4.35 | 15.89 | 3.12 | 176.01 | 17.09 | 4.03 | 167.88 | 39.09 | 12.12 |
| 4 | Activated $CD8^+$ memory T cells | $CD8^+$ $CD45RO^+$ $HLA\text{-}DR^{high}$ | 3.39 | 30.9 | 4.88 | 2.82 | 0.57 | 120.19 | 47.06 | 1.88 | 202.6 | 58.55 | 15.87 |
| 8 | Naïve $CD4^+$ Tregs | $CD4^+$ $CD127^-$ $CD25^{high}$ $CD45RA^+$ $CD69^+$ $HLA\text{-}DR^+$ $PD\text{-}1^{high}$ $TIGIT^+$ | 53.79 | 11.17 | 3.34 | 136.57 | 9.23 | 13.93 | 95.34 | 1.15 | 140.73 | 168.64 | 24.83 |
| 9 | Naïve $CD4^+$ T cells | $CD4^+$ $CD127^+$ $CD45RA^+$ $CCR7^+$ $HLA\text{-}DR^-$ | 83.3 | 7.84 | 32.6 | 1.4 | 197.0 | 1.09 | 0.52 | 73.64 | 5.37 | 0.92 | 0.88 |
| 13 | Memory $CD4^+$ T cells with check point inhibitor expression | $CD4^+$ $CD45RO^+$ $HLA\text{-}DR^-$ $PD\text{-}1^+$ $TIGIT^+$ | 96.03 | 5.67 | 5.06 | 1.31 | 1.8 | 107.91 | 25.32 | 2.57 | 5.64 | 114.61 | 28.09 |
| 12 | Memory $CD8^+$ T cells with check point inhibitor expression | $CD8^+$ $CD127^+$ $CD45RO^+$ $HLA\text{-}DR^+$ $PD\text{-}1^+$ $TIGIT^+$ | 6.18 | 201.12 | 19.84 | 0.94 | 3.16 | 158.06 | 18.88 | 5.44 | 36.06 | 52.06 | 17.99 |

**Table S8. Biochemical Affinity of Clone 52B8 Variants for Human ILT3-His Protein**

| | n | $k_a$ ($M^{-1}s^{-1}$) | $k_d$ ($s^{-1}$) | $K_D$ (nM) |
|---|---|---|---|---|
| Chimeric 52B8 | 4 | 1.7E+06 | 7.9E-04 | 0.46 ± 0.01 |
| Humanized 52B8 | 6 | 1.6E+06 | 1.4E-03 | 0.90 ± 0.11 |
